# Peripheral Nerve Stimulation Optimized Pulses for EPI (POPE) allows high-resolution fMRI

**DOI:** 10.64898/2026.07.22.739360

**Authors:** Renzo (Laurentius) Huber, Dominik Rattenbacher, Bastien Guerin, Haotian Hong, Alessandra Pizzuti, Omer Faruk Gulban, Tina (Wei-Ching) Lo, Azma Mareyam, Kyle Droppa, Jinting Yao, Cole Analoro, Paul Wighton, David Feinberg, Lawrence L. Wald, Rüdiger Stirnberg

## Abstract

**Purpose:** Recent improvements in MRI gradient design and amplifiers, advanced MRI scanners are now routinely utilizing slew rates of several hundred T/m/s. However, full gradient performance cannot be exploited in high-resolution EPI due to peripheral nerve stimulation (PNS) limits. We aim to characterize and mitigate these PNS constraints using a simple sequence modification: PNS-optimized EPI gradient pulse shapes.

**Methods:** PNS-Optimized Pulses for EPI (POPE): we selectively reduce the slew rate of gradient pulses at periods of high predicted PNS spikes, while leaving the rest of the waveform unchanged. PNS sensation was evaluated.

**Results:** POPE allows 7%-35% faster imaging of EPI protocols resolutions of 1mm-0.3mm resolutions without exceeding predicted PNS. With such improvements, POPE allows robust 0.3 mm isotropic fMRI protocols that would have exceeded safety limits without it.

**Conclusion:** POPE facilitates locally precise fMRI activation mapping on clinical 7T scanners at spatial resolution that were previously unattainable due to PNS limitations.

## Purpose

Recent improvements in magnetic resonance imaging (MRI) gradient design and amplifier technology mean that advanced ultra high field (UHF) MRI scanners are now routinely equipped with high performance gradient systems utilizing slew rates of up to 250T/m/s for clinical scanners (7T Terra.X) and 900T/m/s for research scanners (Impulse 7T)^1^. Such high slew rates are particularly advantageous for high-resolution echo planar imaging (EPI) protocols in mesoscale functional MRI (fMRI) applications to address questions on intracortical vascular physiology and cortical laminar microcircuits^1,2^. However, with increasing gradient performance, high-resolution protocols are becoming increasingly limited by peripheral nerve stimulation (PNS)^3,4^. This holds true both for conventional clinical gradients and for PNS-optimized head gradient hardware, despite their differing threshold levels^5^. For safe operation on MRI scanners, PNS is usually estimated in real time as a function of used slew rates accumulating over time. Thus, in Echo-Planar Imaging (EPI) sequences for fMRI protocols, PNS thresholds are commonly exceeded at the end of each EPI ramp period and need to be mitigated by costly reductions of the sequence performance such as the protocols’ overall slew rates (gradient mode) or gradient magnitude (bandwidth) Fig. 1.

**Fig. 1:**
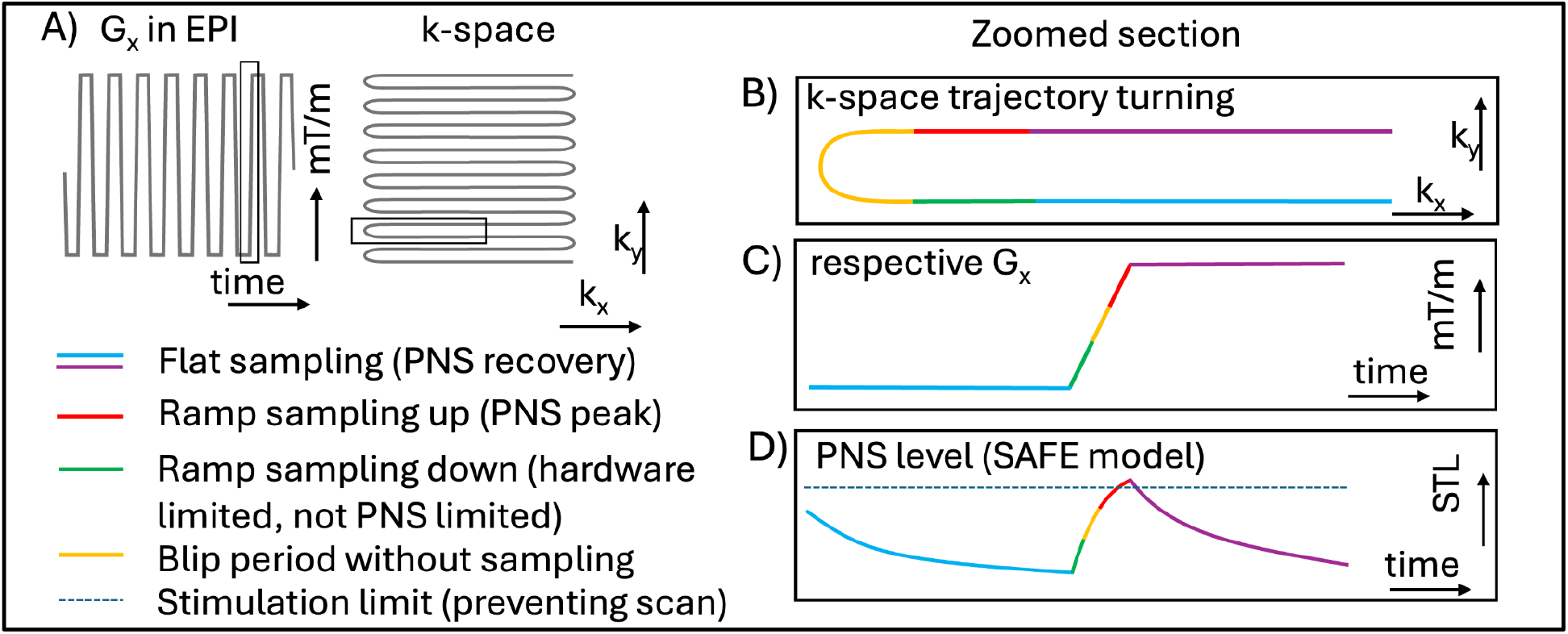
Schematic description of the EPI pulse train with corresponding k-space and respective PNS level. PNS is caused by the slewing while the trajectory is turning directions (zoomed sections of **A** shown in **B-D)**. This slew event includes sampling periods of two independent k-space lines (green and red) and a non-sampling period during the k_y_-blips (yellow). For conventional EPI sequences, these three periods are on the same slew-event. This is not needed however. The individual sub-epochs can be independently modified without violating the linearity requirements of commonly employed ramp-sampling regridding image reconstruction methods. Panel **D)** shows how the temporal rise of the PNS level (mimicking the accumulation of nerve ions) during slew events results in an inverse-exponential shape that peaks at the rising initial corners of EPI flat tops, exceeding the stimulation limit and preventing the scan to be safely executed on the scanner. In POPE, we aim to modify the slew rates along gradient rise, to reduce the PNS peak exclusively in the red period.

Here, we seek to evaluate and optimize EPI pulse shapes for their predicted PNS level using the SAFE model threshold estimation. Previous and parallel studies have investigated pulse shapes to reduce PNS by means of several pulse shape approaches. These include symmetrically cutting off the edges of trapezoidal EPI pulses, using freely curved gradient pulses in diffusion encoding, sign-asymmetric slew rates, and pre-excitation targeting of the potassium system (PRE-TAPS) pulses^6-15^. Here, we are building on this work and propose an EPI-optimized wave form that allows linear “Cartesian” ramp-sampling for vendor-integrated EPI image reconstruction, while exploiting the flexibility of having inherent breaks from data sampling during slew periods at EPI reversals (yellow in Fig. 1B-1C). We focus the EPI pulse shape optimization on the specific case of the PNS SAFE model as utilized on SIEMENS scanners. We aim to implement, validate, test, and disseminate this approach for the application case that we consider severely limited by PNS: high-resolution fMRI.

## Theory and Methods

PNS in human MRI arises from the induced electric fields (E-fields) that accompany the rapidly changing magnetic fields (dB/dt) of switching MRI gradients; when the induced E-field depolarizes a peripheral nerve beyond its excitation threshold, a stimulation is perceived. Nerve excitation follows a strength–duration relationship: the threshold depends not on dB/dt alone, but on how the stimulus accumulates over the nerve’s characteristic integration time. Brief pulses therefore require a higher dB/dt to reach threshold, whereas sustained pulses stimulate at progressively lower dB/dt. The Siemens scanners emulate this temporal integration with the SAFE model (Stimulation Approximation by Filtering and Evaluation^16^), which convolves the gradient slew rate with a set of exponential filters and sums the rectified, weighted outputs into a predicted PNS level that must stay below a coil-specific limit. A publicly accessible implementation of the model is available on GitHub^17^. While the SAFE model is empirical, it was inspired by the physiological behavior of the temporal excitability of nerve cells. The SAFE model is based on a temporal filter of the slew rate with three time constants per gradient axis, which are typically in the range of *τ*1=150-250 *µ*s, *τ*3=0.75-0.85 ms, and *τ*2=12 ms. *τ*3 is usually the most critical time constant for EPI.

For high-resolution EPI protocols with typical echo spacings of 0.6-2.5 ms (Fig. 1A), PNS events are typically sparse and limited to short spikes at the end of intense slew periods of rising corners of EPI trapezoidal gradient pulses (Fig. 1D). On typical modern clinical 7T scanners with body gradients (Terra.X with W60 gradient amplifiers), these PNS spikes are restricting the hardware achievable performance of the gradients (red in Fig. 1D).

Here, we propose to optimize the EPI gradient shape while considering that the temporal dynamics of the PNS resembles an inverse exponential shape (Fig. 1D). This approach takes advantage of flexibility that the “ramp down” period of a negative EPI pulse (green in Fig. 1B-C) does not need to have the same slew rate as the “ramp up” period of the subsequent EPI pulse (red in Fig. 1B-C). This flexibility allows tailored slew rate throttling, solely at the sparse time points, when PNS is exceeded. We propose to find an optimal balance of the slew rate of the green and red segments to reduce the estimated PNS level without changing the total rise time. This approach is termed PNS-optimized Pulses for EPI (POPE).

## Simulation study

To clarify the conceptional working principle of POPE, we simulated a typical 0.8 mm EPI with echo spacings of 1 ms, and slew rates of 200 mT/m, G_max_ of 42 mT/m, as previously used as a field-wide template imaging protocols at 7T for mesoscale EPI^18^ and achievable at 83% of all 7T scanners (including “classical” Magnetom 7T, Magnetom 7T plus, Terra, and Terra.X). PNS levels were estimated in the Vendor-provided IDEA framework (Integrated Development Environment for Applications, Siemens, 2010, North Carolina, USA). We used SAFE model parameters of Terra.X (software version XA60A). The parameters are confidential, a list of the approximate ranges is given in Tab. S1. The temporal evolution of the estimated PNS level is shown in Fig. 2A for conventional EPI pulses, and for POPE EPI pulses.

**Fig. 2:**
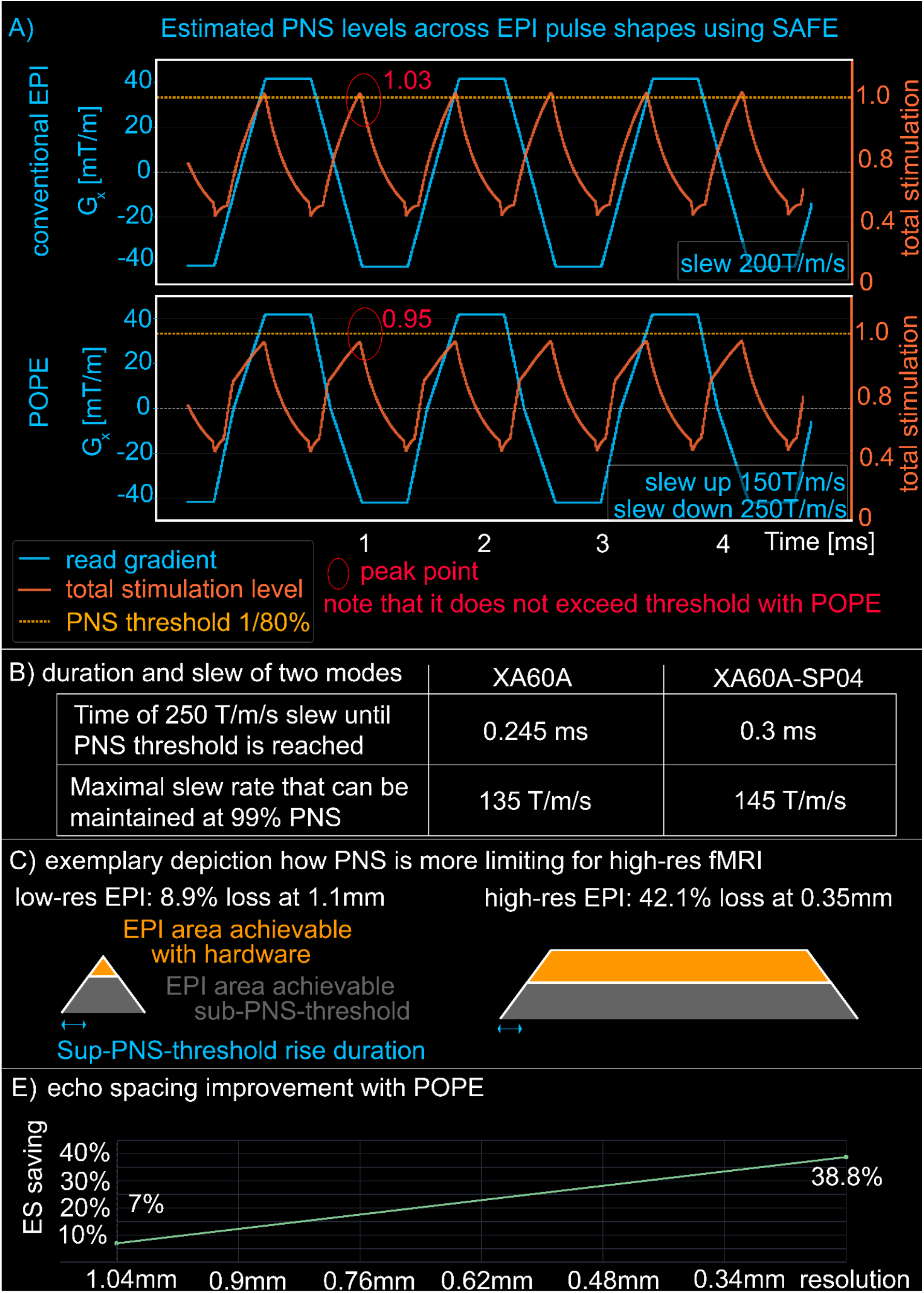
**Panel A)** shows results of the exploratory simulation study of 0.8mm EPI. While conventional EPI exceeds the PNS threshold, balancing the ramp up and ramp down slew rates reduces the peak PNS to subthreshold levels for the same echo spacings. **Panel B)** shows results of the simulation study that determines the duration of sub-threshold PNS with maximal slew and the steady-state slew to maintain 99% of the stimulation level. **Panel C)** depicts two selected example EPI pulse shapes across two resolutions and how they are limited by PNS thresholds. This is to visualize why PNS is having larger effects on high-resolution protocols compared to low-resolution protocols. For low-res EPI with more relative ramp sampling and less time spent on the flat top, PNS-limiting slew durations do not have large effects on the resulting gradient moment. For high-res EPI with less relative ramp sampling and more time spent on the flat top, PNS-limiting slew durations have significant effects on the resulting gradient moment. **Panel D)** Depicts simulation results of the relative improvement in echo spacing that POPE can provide across resolutions. POPE is particularly beneficial at higher resolutions. Insets refer to pulse shape differences at selected edge cases.

Based on the considerations on EPI flexibility (Fig. 1), and the conceptual simulations (Fig. 2.) we decided to implement POPE assuming a two-state model:

- Hardware limited periods: Below the PNS threshold, the maximal slew rate can be used until 99% of the predicted PNS threshold is reached.
- PNS limited periods: As soon as the predicted PNS level is at 99%, a lower “steady-state” slew rate must be employed that results in a constant PNS level. The resulting PNS level plateaus can then be maintained indefinitely.

This simplification holds because typical EPI protocols with echo spacings of 0.5–1.5 ms and whole-body gradient specifications of 150–250 mT/m/s slew rate and ±25–45 mT/m G_max_ (giving slew times of 0.5–1 ms) are dominated by a single SAFE time constant, *τ*_3_ = 0.75–0.85 ms. For other applications, a full multi-exponential gradient wave form with multiple time scales might be needed (e.g. see Hannum 2026 for PNS optimized diffusion^14,15^).

In the simulation study, we estimated the duration that the gradients can use the maximal slew rate of 250 mT/m until 99% of the maximal PNS is reached, and we estimated the “steady state” slew rate that can be maintained at 99% of the predicted PNS level.

## POPE for high-resolution fMRI in vivo

To test the feasibility of POPE for high-resolution fMRI, we implemented POPE into the DZNE 3D-EPI Skipped CAIPI sequence with VASO capabilities^2,19^. See supporting information (Fig. S1) for more implementation details. The sequence branch that includes POPE functionality is publicly available on the SIEMENS “App Store”: C^2^P exchange platform on Teamplay.

Scanning was performed at Magnetom 7T Terra.X XA60A (Siemens Healthineers, Forchheim, Germany) at MGH with W60 Gradient amplifiers supporting G_max_ 135 mT/m (42 mT/m for EPI) and slew of 250 T/m/s (fully utilizable in EPI) upon receiving local IRB approval (2025P002459). N=6 participants were scanned with a custom 64ch 7T coil^20^. Most relevant protocol parameters are: Nominal resolution 0.3 mm iso., matrix 420×420×18, TR 6.2 s, in-plane FOV 130 mm, G_max_ read 41.9 mT/m matched for minimal 2 *µ*s dwell time, 1×3z1 CAIPI undersampling, segmentation 18^19^ = shot-selective CAIPI^21-22^ 6 shots per k_z_-plane, Partial Fourier 77% (EPI factor 18), 3rd order shims disconnected. A full protocol parameter list can be seen on Github: Parameter list: https://github.com/layerfMRI/Sequence_Github/blob/master/POPE/FRISGO_20251008_AZM_AA.pdf.

Vendor reconstruction performed regridding for ramp-up and ramp-down sampling, with GRAPPA kernel fitting in IcePat using dual-polarity averaged ACS data. FRISGO artifact correction was applied using alternating EPI readout polarities^23-24^.

For consistent slab alignment across runs in the presence of subject head motion, we used the AutoRegister framework on a run-by-run basis^25^. In this framework, Low resolution scout sequences are sent to a remote computer for registration, then the AutoAlign matrix is overwritten to update the FOV forsubsequent scans (https://github.com/pwighton/auto-register). For B_0_-shimming, we used the WIP #162^26^. Functional tasks consist of 30 s blocks of flickering checkerboards. 4-5 runs were conducted per participant of 12 min each. The task presentation was presented with PsychoPy^27^ and is available on Github: https://github.com/layerfMRI/Phychopy_git.

Additionally, two participants were repeatedly invited for three scanning sessions using visual and sensory motor stimulation (30 s finger tapping paradigms) to explore the capabilities of POPE to address future questions of multi-scale BOLD signal mechanisms as a higher sensitivity than single-session data can provide^28^. These multi-session experiments used a larger FOV, resulting in 0.35mm resolution (https://github.com/layerfMRI/Sequence_Github/blob/master/POPE/Multi_session.pdf).

We asked all participants after all sessions, whether they felt PNS.

## Results

Simulation results provided quantitative predictions for how long the maximal slew can be used until the PNS threshold is reached (Fig. 2B). And they provided predictions of the steady-state slew rates that maintain sub-threshold PNS levels (Fig. 2B). The durations of maximal slew are smaller than the hardware limitations. As such, the hardware can support slewing at 250 mT/m for 1.08 ms across G_min/max_ ± 135 T/m/s. The PNS limited slew rate of 0.24 ms limits the hardware accessible durations to 26% (73% for EPI with G_min/max_ ± 42 T/m/s). The steady state maintainable with slew rate is 135 T/m/s (Fig. 2B), limiting the hardware accessible performance of 250 T/m/s by 54%.

Simulation results in Fig. 2D shows that POPE can safely achieve shorter echo spacing without triggering PNS, compared to conventional trapezoidal EPI pulses. The benefit of POPE is largest at high spatial resolutions when EPI sampling is dominated at the flat top period.

In in-vivo tests, we found that POPE EPI could be used to achieve EPI resolutions of nominal 0.3 mm voxel sizes. Without POPE, these protocols were unachievable within safety limits or suitable echo times. Representative raw data quality can be seen in Fig. 3. While individual time frames are limited by thermal noise, upon averaging 4-5 runs with identical activation timing, the SNR is improved with tSNR values reaching the double-digit regime and activation maps that exhibit spatial patterns of cortical folding. No indications of potential re-gridding artifacts were detected in these high-resolution images.

**Fig. 3:**
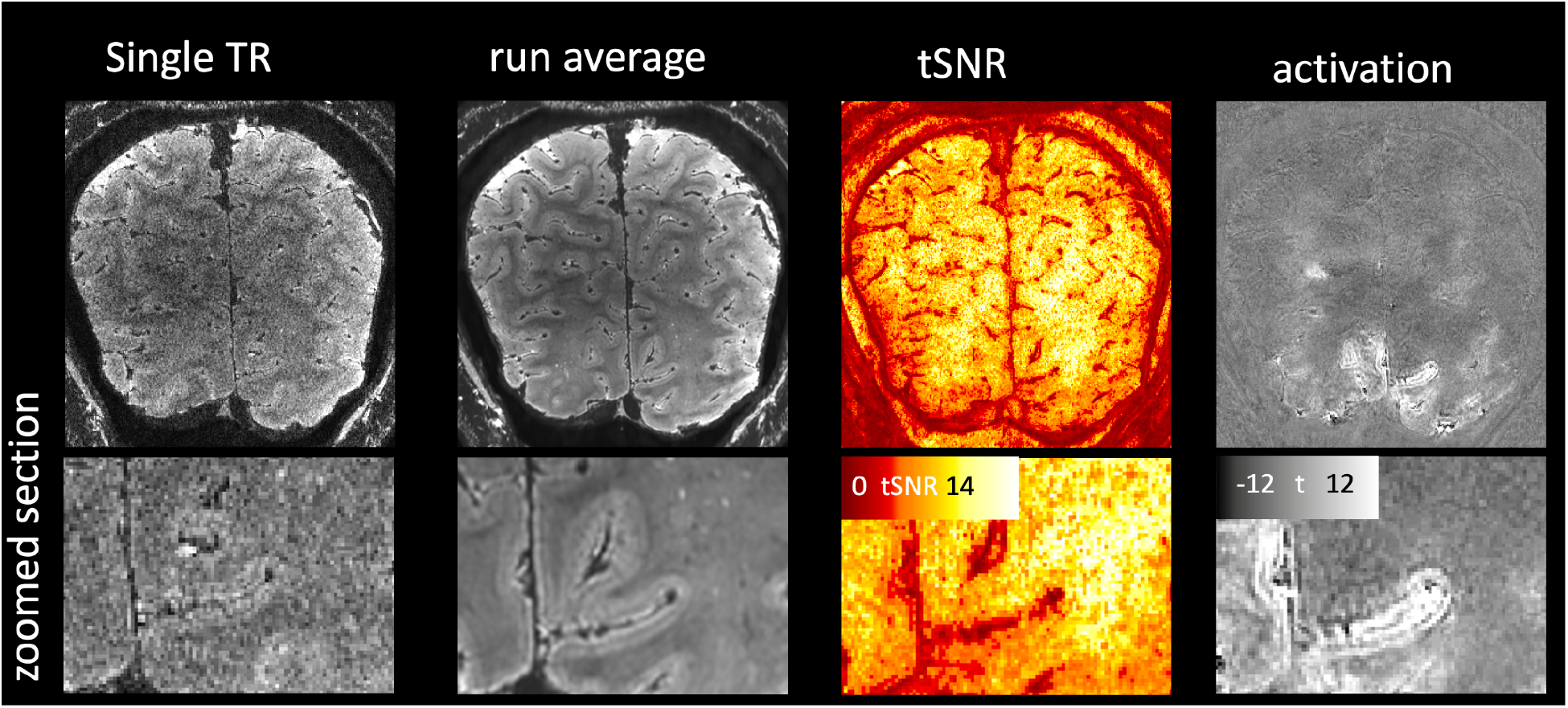
Quality assessment of 0.3 mm isotropic EPI in a representative participant. POPE provides EPI quality that is suitable for fMRI applications. The single TR and run average show the structural fidelity of the acquisition method. The spatial structure of the tSNR map shows that these data are limited by both (physiological noise and thermal noise). E.g. vessels and CSF-GM transition areas have low tSNR values. White matter has higher tSNR than GM - different than the EPI signal. The unthresholded activation map shows that there are no unexpected false positives below the threshold. E.g. there are no indications of edge effects of motion, there are no false positive signals from segmentation related ghosting.

**Fig. 4:**
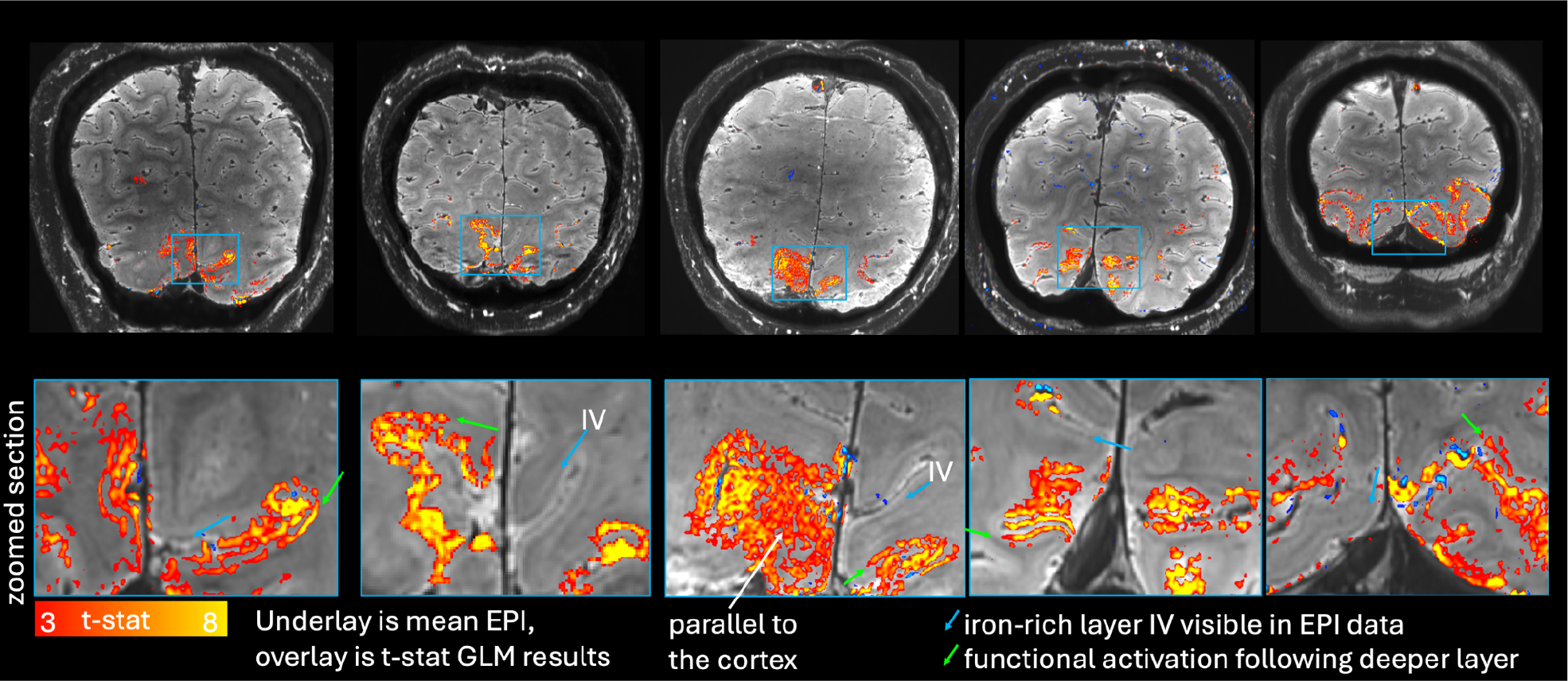
Results across participants. POPE fMRI (0.3 mm isotropic) is stable across all participants. In all participants, fine scale intra-cortical and pial signals are detectable. Note that the underlay grayscale images are directly derived from POPE fMRI time series (average). This is important in addition to the fMRI activity, as we can directly assess the anatomical details such as the cortical layers and mesoscopic vessels.

Fig. 5 depicts multi-session results to highlight the potential of high-resolution POPE to address questions on BOLD signal origins across the spatial scales along the vascular tree. We find that POPE fMRI can spatially distinguish activation from individual intracortical ascending veins (green arrows).

**Fig. 5:**
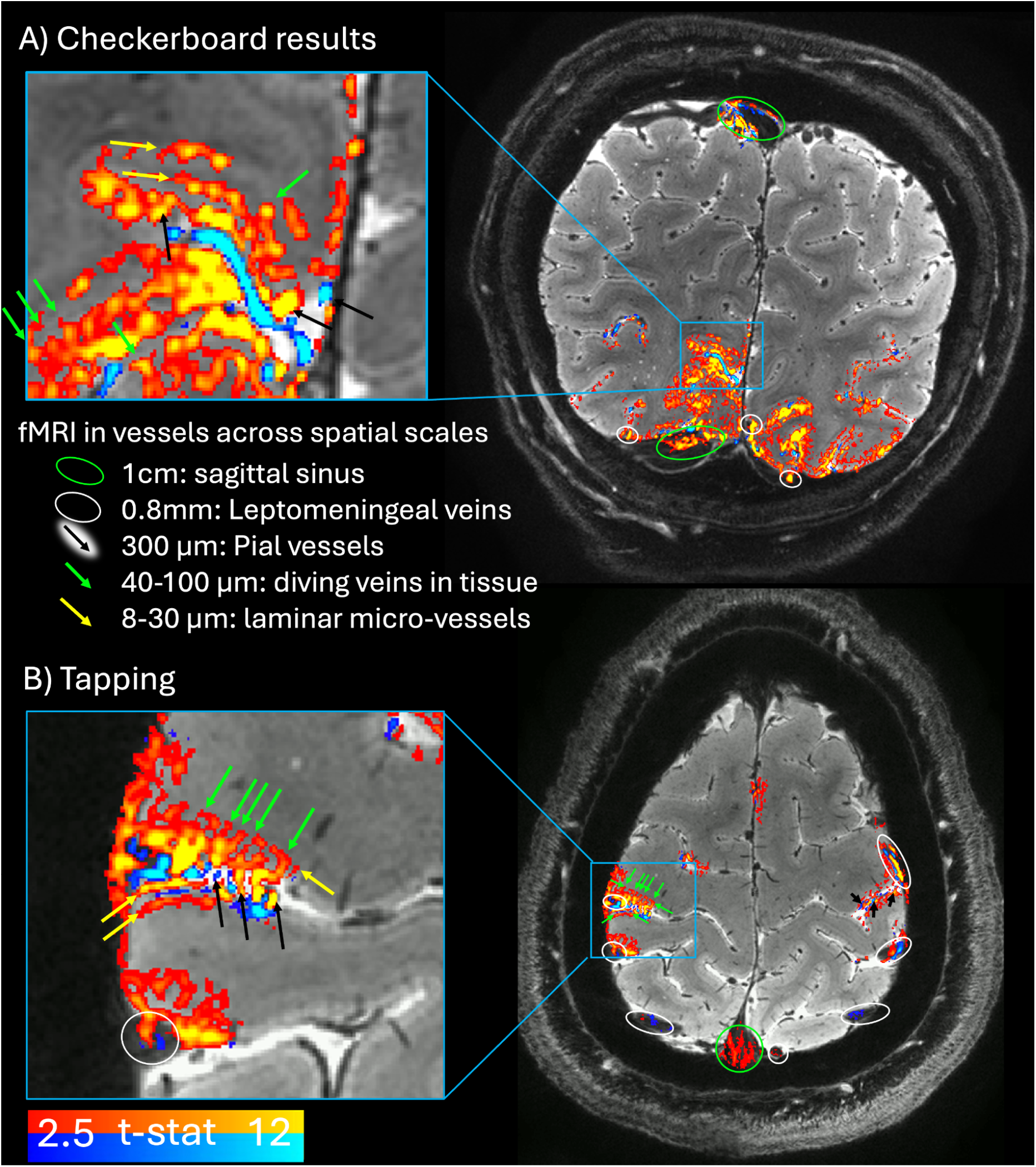
Multi-session results to boost SNR for detection of fMRI signal changes across spatial scales of the vascular tree. Results from the visual (A) and sensorimotor (B) cortices are shown after averaging data across three sessionsData from three sessions were averaged. It can be seen that activation changes can be captured and identified from vessels across 6 orders of magnitudes.

None of the six participants reported the sensation of PNS in any experiments (they were asked after every scan if they felt “something weird”).

During the course of this study, we observed that POPE fMRI protocols allowed such high G_max_ values and duty cycles that the gradient temperature could not be maintained at a sustainable level below critical 60 °C. Above 60 °C, the gradient epoxy is expected to start developing microcracks. This was addressed with 6-minutes gradient cool-down periods between runs, limiting the number of runs per session. The measured temperature evolution of the gradient followed the functions T_heat_(t)=66.5−39.7·exp(−0.244t) and T_cool_ (t)=20.5+39.0·exp(−0.120t), with respective time constants of 1/λ_heat_= 7.14 min and 1/λ_cool_= 5.88 min, respectively measured data in Fig. S3. Based on this finding, we decided to limit G_max_ in future studies below 38 mT/m which keeps the gradient temperature just below the 60°C.

## Discussion and summary

High-resolution fMRI with the latest generation of clinical 7T scanners is significantly limited by induced and estimated PNS, prohibiting the users to embrace modern hardware capabilities. We developed a simple modification in the EPI gradient pulse shape to mitigate PNS solely at the time points of PNS peaks which is compatible with existing vendor reconstructions. This allows 0.3 mm fMRI mapping in 3D. While a conventional 3 mm voxel contains ≈ 700,000 neurons^29^, POPE’s 0.3 mm voxels are specific to ≈ 700 neurons per voxel.

Voxel sizes at these spatial scales are used across a number of research labs to robustly detect fMRI signal changes in parallel efforts, E.g. with advanced artifact mitigation methods FRISGO^28,30^, with high field strengths at 10.5 T^31,32^, and with the Next-Gen 7T head scanner^1^. Here we are achieving these resolutions with “clinical” 7T scanners, which allows wide dissemination. In the first 2 months since the public release of the sequence with POPE capabilities, it has already been downloaded by 20 institutions.

POPE provides benefit specifically when EPI pulses are operating near the full achievable G_max_, with correspondingly long slew periods, as it is typical in high-resolution protocols. Lower resolution fMRI protocols that do not engage the full available range of G_max_ are not PNS-limited on conventional body gradient systems and will not benefit from POPE.

While the chosen empirical feasibility tests of POPE in Fig. 3-5 are focusing on 0.3 mm data, we anticipate that POPE will also be specifically beneficial for 0.8-1 mm fMRI protocols. At this range of resolutions, conventional body gradients of Siemens scanners exhibit an unfortunate interaction of echo spacings at mechanical resonances and PNS. In pilot studies, POPE has shown to provide the flexibility to circumvent mechanical resonances where a relatively modest improvement of PNS level allows a strong improvement in acquisition speed^33^.

In addition to functional imaging, we anticipate POPE to be used in ultra high resolution anatomical imaging targeting below 0.35 mm isotropic voxel with whole human head coverage^35^. By using longer fMRI-style EPI readouts, these anatomical protocols can compress acquisition times from seven minutes to approximately one minute, making POPE essential for motion-sensitive applications. This capability is going to be critical for clinical studies and neuroimaging in special populations, including pediatric and infant cohorts.

We believe that approaches like POPE are beneficial to extend safe execution of high duty-cycle sequences (E.g. EPI and diffusion) on high performance gradient systems as the emerging technology of Cima.X, MAGNUS, Impulse, and Connectome.X.

While no participant reported PNS sensation across any of the experiments, it remains unclear whether the predicted PNS reduction from POPE perfectly matches the behavior of nerves and truly elevates population threshold averages as suggested by the SAFE metric. Because the SAFE model is inspired by the biophysics of nerve excitability as temporal accumulation of ions, we expect that POPE not only reduces the modeled safety metric but also genuinely reduces nerve excitation. Ongoing studies using POPE on the Impulse head scanner aim to confirm this directly.

While POPE allows fMRI protocols with more aggressive resolution, slew rates and G_max_ values, hardware limitations such as gradient heating can become a practical limit of taking full advantage of it. Measured data and model fits of gradient heating are available on Dataverse.

## Data sharing statement

Example 0.3mm data are shared on the Harvard Dataverse: https://doi.org/10.7910/DVN/0A2ZOK. Scan protocols are shared on Github: https://github.com/layerfMRI/Sequence_Github/tree/master/POPE. This repository also contains the sequence dsv files used in Figs.2, S4. The sequence binaries are shared via the Siemens C^2^P exchange platform. No new analysis has been developed for this study. The routine high-resolution analysis scripts used here are available on Github: https://github.com/layerfMRI/repository.

## Acknowledgements

This work is supported by the Center of Mesoscale Mapping (NIH P41EB030006) and the MGH 7T Center. We thank Ana Arsenovic for helping with in-vivo scanning safety regulations/training. We thank Jocelyn Mora for help with IRB protocol submission. We thank Tyler Morgan, Kenny Kan, and Florian Kroh for sharing and testing fMRI protocols and their PNS estimated from other scanners. We thank Ali Aghaeifar for his B_0_-shiming routines in the form of his WIP #172. We thank Handrik Mattern for comments on early presentations of this work leading to the analysis of mechanical resonances shown in Fig. S3.

## Conflict of interest and Diversity statement

The work presented here may be specific to the PNS safety models used in SIEMENS Healthineers’ UHF scanners. This vendor is used in 83% of all human layer-fMRI papers (source: www.layerfmri.com/papers). Dominik Rattenbacher is an employee of SIEMENS Healthineers. Faruk Gulban is an employee of Brain Innovation B.V.. Recent research has identified a bias in citation practices, with papers from women under-cited relative to their representation in the field^34^. In the human layer-fMRI community, average gender citation bias is 84% male, 15% female (https://layerfmri.com/papers/). For this paper’s references, we determined first-author gender (excluding self-citations) and found 70% male, 30% female. We look forward to future work supporting more equitable citation practices.

