## Supplementary tables and for "Peripheral Nerve Stimulation Optimized Pulses for EPI (POPE) allows high-resolution fMRI"

### Supplementary Material to the manuscript: Peripheral Nerve Stimulation Optimized Pulses for EPI (POPE) allows high-resolution fMRI

By Renzo (Laurentius) Huber[1,2], Dominik Rattenbacher[2], Bastien Guerin[1,2], Haotian Hong[1,2], Alessandra Pizzuti[4], Omer Faruk Gulban[4,5], Tina (Wei-Ching) Lo[1,2], Azma Mareyam[1,2], Kyle Droppa[1,2], Jinting Yao[1,2], Cole Analoro[1,2], Paul Wighton[1], David Feinberg[6], Lawrence L. Wald[1,2], Rüdiger Stirnberg[7]

1. Mass General Brigham, 2. Harvard Medical School, 3. SIEMENS Healthineers, Forchheim, Germany, 4. CN, FPN, Maastricht University, Netherlands, 5. Brain Innovation, Maastricht, Netherlands, 6. UC Berkeley, 7. DZNE Bonn, Germany

**Tab S1: Example model parameters used in the simulation study here.**

| Model parameter | Terra.X XA60A |
|---|---|
| Stimulation Limit X | ≈25-35 |
| Stimulation Limit Y | ≈25-35 |
| Stimulation Limit Z | ≈15-25 |
| $\tau_1$ | ≈50-200μs |
| $\tau_2$ | ≈10-20ms |
| $\tau_3$ | ≈0.5-1ms |

*The values refer to approximate ballpark numbers. The exact numbers are proprietary information and cannot be shared publicly. Instructions on how users can see the exact numbers of their gradient system are given here: https://github.com/filip-szczepankiewicz/safe_pns_prediction. From software version XA60A SP04 onwards, the SAFE Model has an updated parameter set. Due to a rebalancing of the gradient axes, more performance on the X gradient is allowed. Nevertheless, the updated parameters fall into the same ballpark ranges as those shown here.*

**Fig. S1: Implementation details of POPE vs. conventional acquisition w/o and with ramp sampling.**

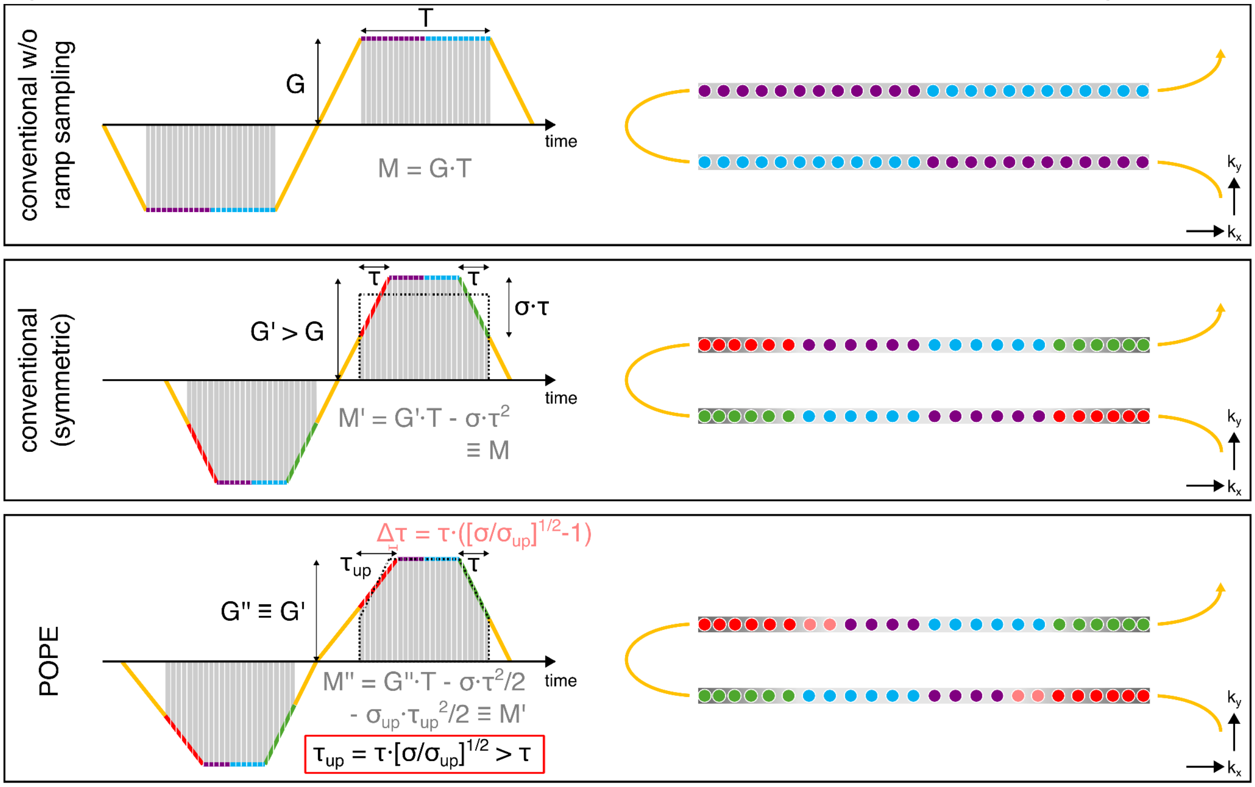


***Fig. S1:*** *For initial feasibility and simulation experiments, we have increased the rise duration with a simple scale factor while leaving everything else unchanged, including ramp sampling. For later functional experiments and public sharing of the sequence, a full POPE implementation has been conducted that includes details of shifted gradient moments in variable ramp sampling. This figure explains this implementation.*
*Left: two gradient lobes and 0th order gradient moments (gray) accumulated during total signal sampling time (T).*
*Right: k-space trajectory and sampling illustrated by colored circles (cf. Fig. 1; reddish: during ramp up, green: during ramp down) with constant or variable sampling density illustrated by grayish background. Conventional ramp sampling (equal time $τ$ on both ramps, e.g. with maximal slew rate $σ$) requires a higher amplitude G'>G than without ramp sampling. Keeping the ramp sampling times unchanged when increasing the rise duration results in a slightly increased 0th order moment accumulated during T, unless the gradient amplitude is reduced accordingly (≤G') .*

*In the present implementation of POPE, we reduce the ramp up slew rate ($σ_{up}$<$σ$) starting from an optimal baseline (symmetric ramp sampling with the highest allowed slew rate $σ$), but we seek to keep the amplitude unchanged (G''≡G') by adding just the right amount of ramp up sampling ($τ_{up}$ = $τ$+$Δτ$) to maintain 0th order moments (M''=M'). This results in $Δτ$ = $τ$(√[$σ$/$σ_{up}$]-1). The total echo spacing ($ESP_{POPE}$) does not increase as much as it would with unchanged ramp sampling times ($ESP_{POPE}$+$Δτ$, if $τ_{up}$=$τ$). Symmetric relaxation of the slew rates for the same amount of PNS reduction typically leads to longer ESP (cf. Fig. 2A). In practice, actual ramp times, slew rates and amplitudes may still vary slightly as the gradient timing needs to be defined on the gradient raster time (10µs). M''≡M'≡M is still maintained by final amplitude downscaling (ensured by adequate rounding of gradient timing).*

**Fig. S2: Measured gradient temperature with EPI utilizing ±42 mT/m $G_{min/max}$.**

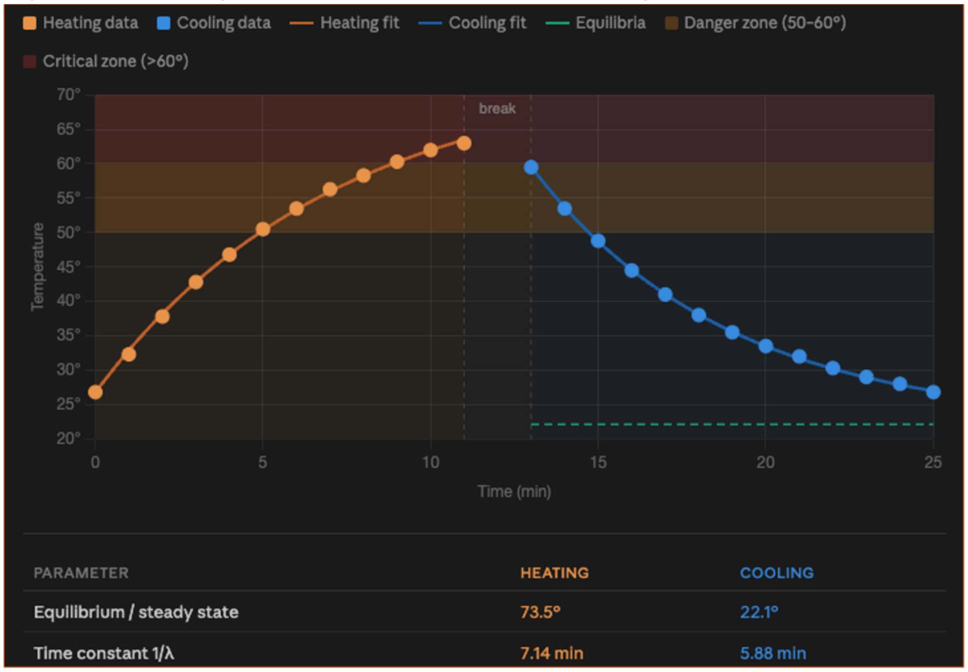


*Measured gradient heating and cool down of the 0.3mm protocol with the gradient model SC72.*

**Fig. S3: temporal spectrum of EPI with and without POPE using identical echo spacing.**

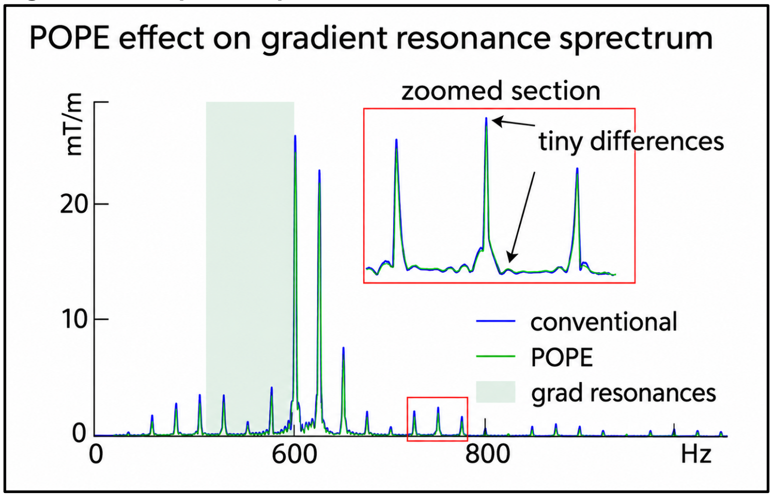


*It can be seen that the altered pulse shape with POPE does not affect mechanical resonances at a significant level. The spectrum refers to the protocol as shown in Fig. 2.*

**Fig S4: Scope measurements of the EPI trajectories with and without POPE.**

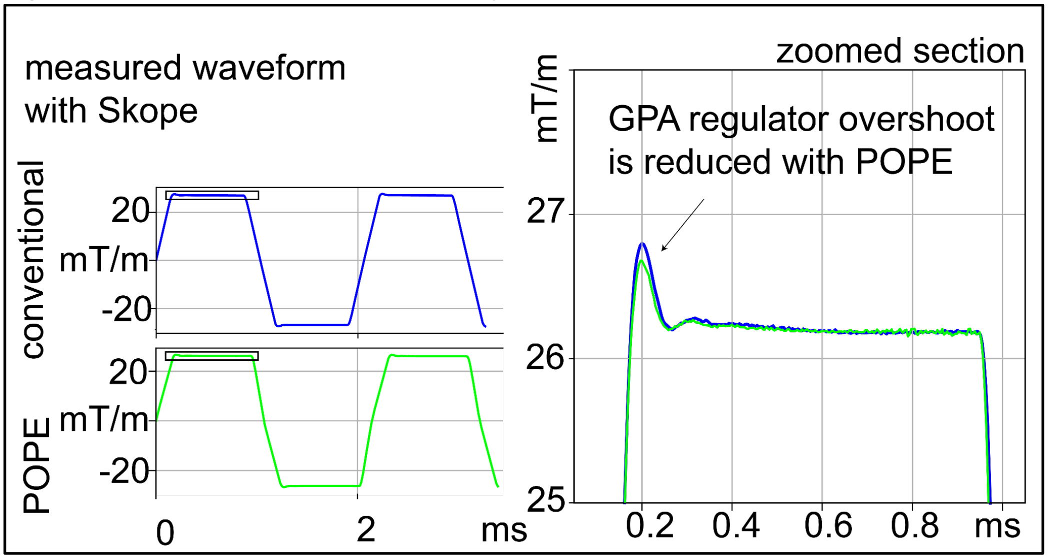


*It can be seen that trajectory imperfections are driven by GPA overshoot across all EPI wave forms. The amount of the overshoot is determined by the slew rate. Thus, with the reduced slew rate of POPE, it also reduces the overshoot (ringing) of the regulators from the gradient amplifiers (GPA). The measured trajectories are referring to a protocol as used in Fig. S3 and 2. The bandwidth, and therefore the readout gradient amplitude, was scaled down to make it executable at the scanner without triggering the PNS watchdog. For the zoomed section plot, the time onsets of the POPE and ANTI-POPE trajectory measurements were shifted (backwards and forwards, respectively) so that the readout flat top times coincide with the conventional trajectory despite varying ramp up times.*
